# The relationship between robustness and evolution

**DOI:** 10.1101/268862

**Authors:** Pengyao Jiang, Martin Kreitman, John Reinitz

## Abstract

Developmental robustness (canalization) is a common attribute of traits in multi-cellular organisms. High robustness ensures the reproducibility of phenotypes in the face of environmental and developmental noise, but it also dampens the expression of genetic mutation, the fuel for adaptive evolution. A reduction in robustness may therefore be adaptive under certain evolutionary scenarios. To better understand how robustness influences phenotypic evolution, and to decipher conditions under which canalization itself evolves, a genetic model was constructed in which phenotype is explicitly represented as a collection of traits, calculated from genotype, and the degree of robustness can be explicitly controlled. The genes were subjected to mutation, altering phenotype and fitness. We then simulated the dynamics of a population evolving under two classes of initial conditions, one in which the population is at a fitness optimum and one in which it is far away. The model is formulated with two robustness parameters in the genotype to phenotype map, controlling robustness over a tight (*γ*) or a broad (*α*) range of values. Within the robustness range determined by *γ*, high robustness results in a equilibrium population fitness closer to the optimal fitness value than low robustness. High robustness should be favored, therefore, under a constant optimal environment. This situation reverses when populations are challenged to evolve to a new phenotype optimum. In this situation, low robustness populations adapt faster than high robustness populations and reach higher equilibrium mean fitness. A larger set of phenotypes are accessable by mutation when robustness is low, in part explaining why low robustness is favored under this condition. A larger range of robustness could be sampled by varying α, revealing a complex relationship between robustness and both the initial rate of phenotypic adaptation as well as the final equilibrium population mean fitness. Intermediate values of α produced a bifurcation in evolutionary trajectories, with some populations remaining at low population mean fitness, and others escaping to achieve high population mean fitness. We then allowed robustness itself to be encoded by a mutable genetic locus that could co-evolve along with the phenotype under selection. Low robustness genotypes are initially favored when adapting to a new optimal phenotype. A high robustness genotype then replaces it, well before maximum fitness is achieved, and moreover appears to prevent further invasion into the population of a low-robustness genotype. This phenomenon was dependent on having tight linkage (and sufficiently low mutation rate) between the robustness locus and the loci encoding phenotype.

## Introduction

Phenotypes display a spectrum of sensitivities to environmental cues. At one extreme are highly plastic traits, for which specific environmental or physiological signals induce alternative optimal phenotypes. At the other extreme are invariant, i.e., canalized or robust, traits for which the influence of the external environment has been attenuated. Natural selection can establish setpoints along this spectrum of environmental sensitivity to optimize populations to specific environmental regimes [Ancel, 1999, Stewart et al., 2012, Ehrenreich and Pfennig, 2015, Palacio-López et al., 2015].

Here we address questions concerning phenotypes that fall at one end of the spectrum— canalized traits—and how they evolve. First brought to our attention by C. H. Waddington [Waddington, 1942], canalization involves not only the attenuation of environmental variability, but also the insensitivity to genetic mutation and developmental noise. Canalization is widely recognized as a general feature in developmental processes, due both to the structural properties of developmental networks as well as to evolution of traits governed by stabilizing selection [Siegal and Bergman, 2002, Barkai and Leibler, 1997, Meiklejohn and Hartl, 2002, Félix and Barkoulas, 2015]. Recent studies investigating gene regulatory networks have also found support for Waddington’s classical idea of canalization being governed by attractor states in the epigenetic landscape [Manu et al., 2009a,b, Moris et al., 2016, Huang, 2012].

The high reproduciblity of a strongly canalized trait attenuates heritable phenotypic variation, thus reducing the rate of adaptation [Fisher, 1930]. Yet canalized traits evolve adaptively over phylogenetic time. Scutellar bristle number in *Drosophila*, for example, is highly canalized [Rendel, 1959, Wheeler, 1987]; but also differs characteristically within family Drosophilidae [Wheeler, 1987], making it a useful taxonomic feature in species classification. Under what circumstances, then, can canalized traits evolve? Early conceptual models invoke gene-environmental interactions upon specific environmental stimuli to release phenotypic variation on which natural selection can act [Waddington, 1942, 1953, Simpson, 1953]. This process involves environmental factors interacting with existing genetic variation to create novel phenotypes [Scharloo, 1991, Gibson and Hogness, 1996, Gibson and Dworkin, 2004].

Mutation can also release otherwise suppressed phenotypic variation. The number of vibrissae is typically nineteen in the house mouse, but in *Tabby* mutants this number can vary [Dun and Fraser, 1958]. Similarly in *Drosophila*, the number of scutellar bristles varies in a *scute* mutant but not wildtype [Rendel, 1965]. Release of phenotypic variation is not restricted to mutation in genetic components regulating a specific phenotype. There may also be genetic factors that act as global regulators of robustness, sometimes called “evolutionary capacitors”. Rutherford and Lindquist [1998] proposed that Hsp90, a chaperone involved in protein folding, is one such factor, buffering phenotypic variation under non-stressed environmental conditions but releasing this variation at opportune times in response to specific environmental stressors [Gibson, 2009, Masel, 2005, Paaby and Rockman, 2014, Masel and Siegal, 2009]. Additional regulators have been found in a genetic screen for them in yeast [Levy and Siegal, 2008]. But evidence for the existence of genetic capacitors that contribute to adaptation remains under debate [Masel and Siegal, 2009, Meiklejohn and Hartl, 2002, Levy and Siegal, 2008, Masel, 2005]. The strongest support for this hypothesis comes from a study that found directionality of eye size change upon Hsp90 induction in the cavefish, a phenotype that is otherwise stabilized by Hsp90 in the ancestral surface populations [Rohner et al., 2013]. More recently, however, Geiler-Samerotte et al. [2016] found a destabilizing effect of Hsp90 on *de novo* mutations, indicating that the buffering effect of Hsp90 on standing population variation involves a selective sieve against destabilized mutants.

The largest gap in our understanding of how canalized traits evolve, however, does not revolve around uncertainly about the specific molecular mechanisms of canalization that contribute to adaptation (such as Hsp90). Rather, it revolves around theoretical issues: the degree of robustness that can be maintained in the population under stabilizing selection, and the extent to which a lowering of robustness is required when adapting to a novel environment. Here we summarize key findings about robustness against mutation.

A variety of empirical, mathematical and computational approaches explore the relationship between robustness and the rate of adaptation, but offer little consensus [Bloom et al., 2006, Hayden et al., 2011, Cuypers et al., 2017, Wagner, 2008, 2012, Draghi et al., 2010, Mayer and Hansen, 2017]. Many studies invoke a conceptual model of a genotype network (originally referred to as neutral network) [Wagner, 2008, 2012, Draghi et al., 2010], which makes the assumption that one optimal phenotype is maintained in the population; all alternative phenotypes are assumed lethal in the current environment. Different genotypes producing the same optimal phenotype that can be connected by a single (one step) mutation form a neutral network. In the face of a new environment in which the current phenotype is no longer optimal, adaptation is achieved by positive selection acting on new mutant genotypes that are one step away from particular (*i.e*., not all) genotypes in the neutral network. This widely studied neutral network model, therefore, invokes epistasis between new mutations and pre-existing alleles in the neutral network to create novel phenotypes. Evolvability here is defined as the probability of a mutation expressing an alternative phenotype.

At its two logical extremes, the neutral network model produces qualitatively different predictions about relationship between robustness and evolvability. At one extreme— complete robustness—wherein the phenotypic effect of mutation is suppressed and thus neutral, the neutral network expands to the full genotypic space and evolvability, by definition, is zero. In this case, evolvability increases when robustness is reduced. At the other extreme— no robustness—as exemplified in [Wagner, 2012], all mutations express an alternative phenotype (assumed to be lethal in the current environment), so the population can only maintain the current genotype. Increasing evolvability results from increasing robustness from this extreme. This is achieved by expanding the neutral genotype network, thereby expanding it’s mutational accessibility to additional phenotypes. But situations close to this extreme may not be evolutionary relevant, since most mutations will produce a non-favorable phenotype and genetic load will be high [Crow, 1963]. A mathematical model that captures the essential features of the neutral genotype network finds that lower robustness can promote evolvability when only a subset of possible phenotypes are mutationally accessible to each neutral genotype [Draghi et al., 2010].

There are other scenarios with different sets of assumptions where greater robustness can promote evolvability. Bloom et al. [2006], for example, show that a protein selected for higher stability while maintaining its native ligand-binding function accumulates a greater number of functionally neutral variants that can potentially bind alternative ligands. Robustness here is defined as maintaining the native function, and evolvability refers to a new dimension of phenotype that was not previously under selection. Under this scenario, higher robustness allows this cryptic genotypic variation to build up, thus promoting evolvability in the new environment. The key difference of this type of model with the previously described genotype network is that there is no selective cost to generating alternative phenotypes here, and the larger genotype network can give rise to a greater number of alternative phenotypes in the new phenotypic space. Experiments or mathematical models in this category include maintaining structural stability when evolving new functions in RNA and protein [Bloom et al., 2006, Hayden et al., 2011], or maintaining homeostasis while evolving new functions [Cuypers et al., 2017]. This type of model is in essence the same as the ones that involve gene-environment interaction to release phenotypic variation under environmental induction, where the same genotype is now evaluated by a hidden dimension of phenotype that is only present under the new environment [Gibson and Dworkin, 2004]. However, although the hidden dimension of phenotype can accumulate variants that may potentially promote the speed of adaptation, as the hidden dimension gets large, the positive relationship may not be guaranteed. The complexity of the relationship of robustness to evolvability builds up further when considering a variable genotype space, with gene duplication and deletion [Crombach and Hogeweg, 2008], or with particular hierarchical structures of gene regulatory networks for robustness (such as a global regulator) [Grigoriev et al., 2014].

Recognizing the complexity of existing models and their differing assumptions, we describe here a tractable genetic model of a multidimensional phenotype to address questions regarding robustness and evolvability. The model employs an explicit genotype to phenotype map, fitness function, and has evolvable control of robustness. For simplicity, genotypes and phenotypes are encoded using Boolean vectors. Our model: (1) draws on established principles of gene regulatory control of traits (we hypothesize gene regulatory networks that determine trait values); (2) specifies a genotype to phenotype map (multiple genotypes map to multiple traits) that is (3) calculated from gene expression states (ON/OFF) and the regulatory activities of the proteins they produce (activating/repressing); (4) has robustness parameters that control the sensitivity of each mutation; (5) allows mutational inputs to the robustness parameter, enabling (5) dynamical analysis of evolution of both traits and robustness simultaneously via a standard Wright-Fisher population genetic model with selection. We employ the model to explore the relationship between mutational robustness and evolvability for populations at the phenotypic optimal and for populations that are challenged to evolve towards a novel phenotype.

## Materials and methods

### A. Model description

The model consists of two parts: the genotype to phenotype map, and the phenotype to fitness map.

#### a. Genotype to phenotype map

We represent the genotype to phenotype map by a one-layer perceptron which maps genotype ***υ*** to phenotype ***z*** using the equation

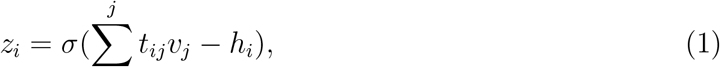

where *z*_*i*_ ∈ {0, 1} is the *i*th component of *z* and *v*_*j*_ ∈ {0, 1} is the *j*th component of ***υ***. Each component *t*_*ij*_ ∈ {−*γ*, 0, *γ*} of ***T*** represents the action of terminal differentiation genes which produce a trait. *h*_*i*_, a real number, is the *i*th component of the threshold ***h***. *σ*(*y*) is a step function such that *σ*(*y*) = 0 iff *y* ≤ 0, and 1 otherwise. ***υ*** has *L* components while *z* and ***h*** have *n* components. In simulations, we assign each element in ***υ*** a value of zero or one randomly in such a way as to ensure that the proportion of ones in ***υ*** is *a*. Each *υ*_*i*_ represents the gene expression state (ON/OFF) of gene *i*. If *υ*_*i*_ equals 0, the gene is OFF (not expressed) and it cannot contribute to any trait. Alternatively, when *υ*_*i*_ equals 1, the gene is ON (expressed), in which case it can have three possible roles in determining a phenotype: activator (*υ*_5_ for phenotype 2 [right panel] in Figure 1A); repressor (*υ*_5_ for phenotype 1 [left panel] in Figure 1A); no functional role (*υ*_6_ for phenotype 1 [left panel] in Figure1). The probability that each *t*_*ij*_ has a non-zero value is given by the parameter *c*, so that *Pr*(*t*_*ij*_ = *γ*) = *Pr*(*t*_*ij*_ = −*γ*) = *c*/2, and *Pr*(*t*_*ij*_ = 0) = 1 − *c*. It is evident that in order for trait *i* to be present (*z*_*i*_ = 1), ∑^*j*^ *t*_*ij*_ *υ*_*j*_ must be greater than the threshold value *h*_*i*_. value *h*_*i*_.

**Figure 1:**
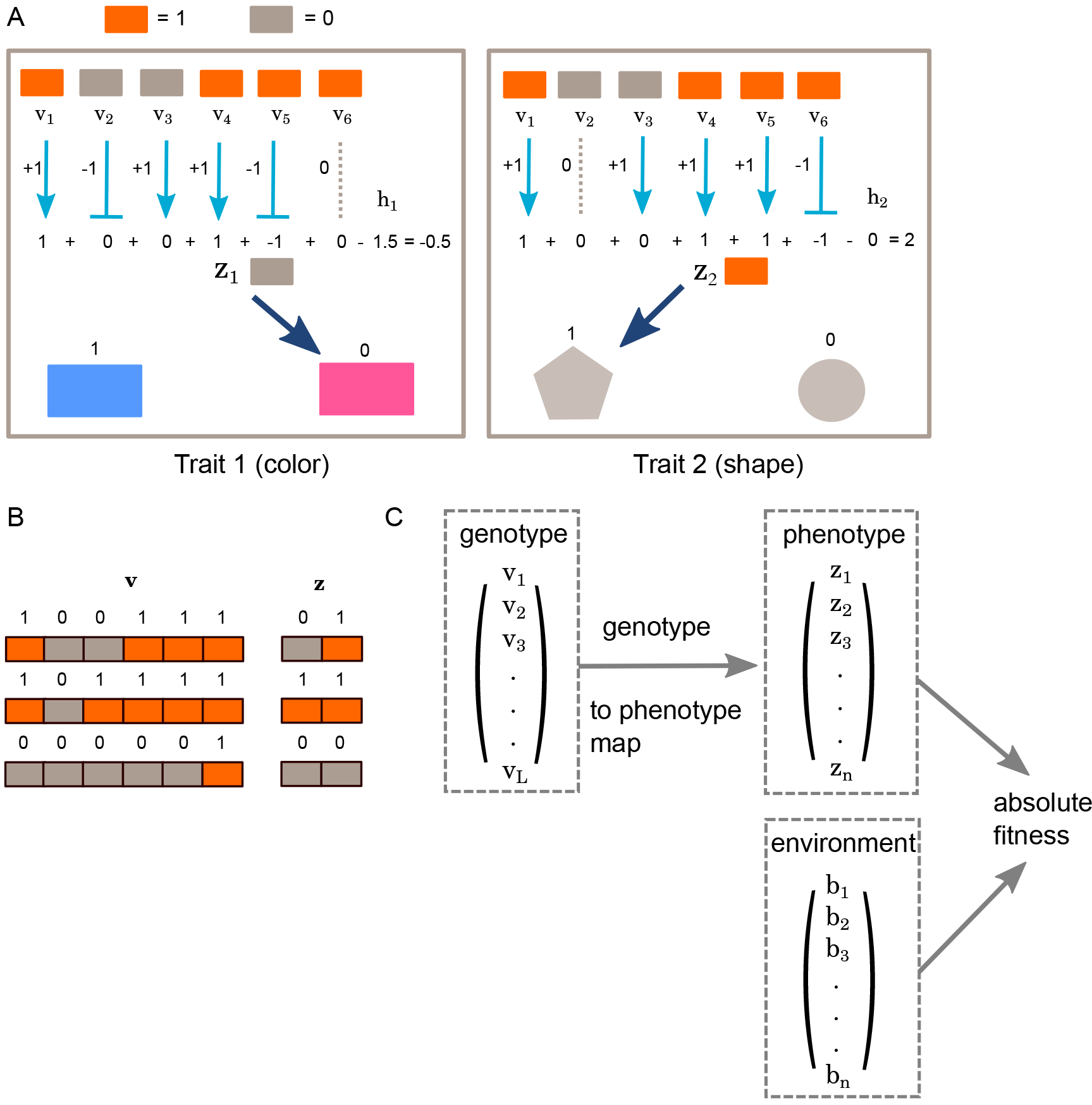
Schematic example and overview of the model. A. Schematic figure showing an example of two traits determined by the genotype
to phenotype map within one individual, from equation (1). Trait *z*_1_ determines color, and *z*_2_ determines the shape. In this example, the number of genes regulating the phenotype is 6, *γ* is set to 1. Interactions of each gene on each phenotype are marked by arrows. Activation is represented by arrows with a triangle end, while repression is represented by a bar end. Grey dashed line means no effect. Under the current configuration, *z*_1_ = 0 and *z*_2_ = 1. B. Examples of different genotype configurations that give rise to different phenotype, given the genotype to phenotype map in A. Orange color indicates “1”, and gray indicates “0” for each genotype or phenotype element. Note that under this particular genotype to phenotype map, phenotype vector *z* cannot take value of (1, 0) by any configuration in genotype, showing the natural constraint by our model. C. Overview of the full model. Genotype is ***υ***, phenotype is ***z***. First part is a genotype to phenotype map, and the second part is phenotype to fitness map. Selection is on phenotype, evaluated by ***b***, representing environment.

#### b. Calculation of robustness

Consider the argument to *σ* in (1), given by

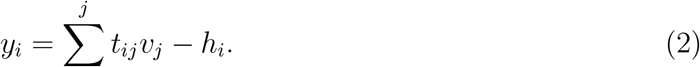

We define the proportion of ***υ*** which is 1 to be *a*. We let the elements *t*_*ij*_ in the *T* matrix to be i.i.d, and *h*_*i*_ is drawn from a Normal distribution 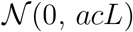, which is independent of *γ*. We can derive the probability of change in *z*_*i*_ when *υ*_*j*_ mutates.

Before mutation, when *L* is large, using the central limit theorem,

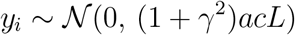

When *υ*_*k*_ mutates, the probability of flipping bits in *z*_*i*_ is given by

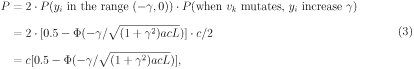

where Φ(*x*) represents the CDF of the standard normal distribution. This argument shows that *γ* is a parameter that controls robustness.

An alternative way to control robustness can be obtained by rescaling *h*_*i*_. We rewrite equation (2) as

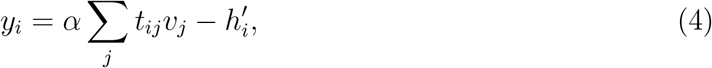

where 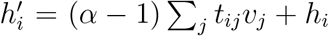. Then the probability that a mutation will change a traitis given by

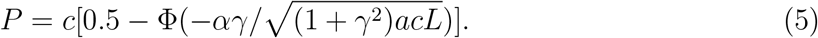

Thus, at fixed *γ*, robustness can also be controlled by varying *α*.

#### c. Phenotype to fitness map

Fitness is a function of phenotype *z* and environment ***b***, where the components *z*_*i*_ and *b*_*i*_ are ∈ 0, 1. The dependence of fitness on ***z*** and ***b*** could be quite complex, but in the present application we consider the environment to specify a trivially explicit optimum phenotype, which denotes the most fit state for each trait in such an environment. An individual’s absolute fitness *w* is defined as the proportion of *n* traits in a phenotype *z* that match those of an optimal phenotype ***b***, so that

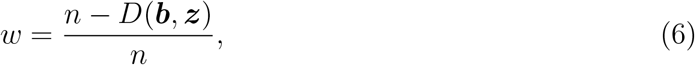

where *D*(***b***, *z*) is the Hamming distance between the two vectors.

### B. Forward evolutionary simulation

A haploid Wright-Fisher forward population simulation was employed to simulate evolution using the parameter values given in Table 2. The threshold *h*_*i*_ is drawn from a Normal distribution 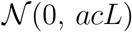.

In each generation, *μLN* mutations are introduced into the population. Each mutation corresponds to flipping a randomly selected bit in ***υ***. The phenotype is calculated for each new genotype, the fitness is calculated from (6), and the next generation is sampled proportionately to the fitness.

These simulations are performed under three enviromental regimes. For E_opt_, the “optimal environment,” the environment ***b*** is set to be the initial phenotype ***z*** in the population. A “new environment” E_new_ is represented by two possible types of environment. In the first, ***b***\* is set to a random vector. Note that there is no guarantee that ***υ*** can generate a maximal *w* in ***b***\* given fixed *t*_*ij*_. We therefore define a second type of new environment, ***b***\*\*, in which optimal fitness is guaranteed to be reachable within the genotypic space. ***b***\*\* is initialized by drawing a random genotype vector and selecting a random genotype-phenotype map, setting ***b***\*\* to be optimal for this map as was done with E_opt_, but using a re-randomized genotype vector to start the simulation.

For simulations with a fixed level of robustness, genes in genotype ***υ*** are fully linked, with mutation rate *μ* per gene per generation. For simulations with evolving levels of robustness we add a robustness locus ***r*** which mutates at rate *μ*_*r*_. Eight levels of robustness are possible, represented by three bits (Table S1). Following mutation, the robustness locus is evaluated first to determine the appropriate *α* value for that individual, which is then applied to the genotype ***υ*** to calculate phenotype. The genes in ***υ*** remain completely linked, but recombination is possible between the genotype ***υ*** and the robustness locus ***r***. The recombination rate ***r*** can range from 0 to 0.5. This was implemented by exchanging robustness loci in Poisson (2*rN*) haploid pairs.

The evolutionary simulation was implemented under the C++ template framework of the fwdpp model [Thornton, 2014] with modification. The Eigen package [Guennebaud et al., 2010] was used to calculate genotype to phenotype map.

Code is available on github (https://github.com/pyjiang/ct_fwdpp).

## Results

Our genetic model attempts to capture essential features of a multidimensional phenotype consisting of a large number of distinct traits controlled by multi-locus pleiotropic gene regulation. For simplicity, we consider a haploid organism with complete linkage of genes. With our main focus being the evolutionary dynamics of robustness, we simplify developmental dynamics by course-graining it into a fixed genotype-to-phenotype map. Genotype is represented by a Boolean vector, ***υ***, whose *L* genes are each set to either an ON (*υ*_*i*_ = 1) or OFF (*υ*_*i*_ = 0) state. The two states can be thought of as representing whether or not a gene is expressed. The phenotype *z* is a collection of *n* traits, also represented by a Boolean vector (Eq. 1). A specific trait in the phenotype might be the presence (*z*_*i*_ = 1) or absence (*z*_*i*_ = 0) of a particular morphological feature, such as a bristle or cell type, or alternatively two distinct states of a trait, such as a yellow or grey body color.

The impact of gene *j* on trait *i* is set by elements *t*_*ij*_ of the two-dimensional ***T*** matrix, with values (*γ*, 0, or −*γ*). *t*_*ij*_ = 0 means that gene *j* is not involved in regulating trait *i*. Positive and negative values of *γ* are interpreted as representing either an activating or repressing contribution of gene *j* on trait *i*. *γ*, in this case, represents the strength of each activation or repression. The genotype-to-phenotype mapping function also has a threshold variable *h*_*i*_ for trait *i*, whose value must be exceeded by the sum of the activation and repression strengths of the genes affecting the trait in order for that trait to be ON. The detailed description of the model is in Materials and Methods and the list of variables is shown in Table 1.

**Table 1:**
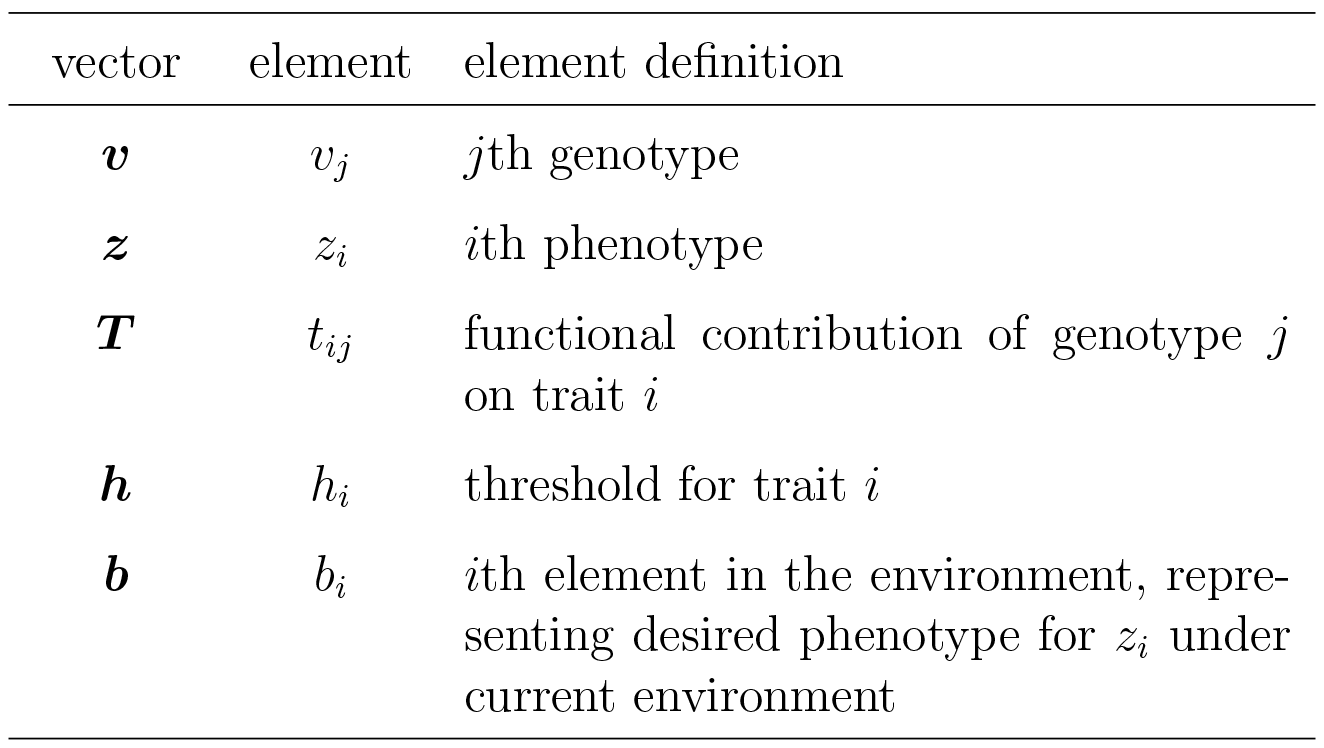
List of variables

| vector | element | element definition |
| --- | --- | --- |
| $\mathbf{v}$ | $v_j$ | $j$th genotype |
| $\mathbf{z}$ | $z_i$ | $i$th phenotype |
| $\mathbf{T}$ | $t_{ij}$ | functional contribution of genotype $j$ on trait $i$ |
| $\mathbf{h}$ | $h_i$ | threshold for trait $i$ |
| $\mathbf{b}$ | $b_i$ | $i$th element in the environment, representing desired phenotype for $z_i$ under current environment |

Figure 1A shows an example in which six genes regulate two traits. With the given interactions, trait *z*_1_ is OFF and trait *z*_2_ is ON. Only mutation in an individuals genotype is allowed to alter its phenotype; the genotype-to-phenotype map remains the same, and there is no influence of environment on phenotype in the model. In the example, *h*_1_ = 1.5 imposes a greater restriction on the conditions for trait 1 to be ON than does *h*_2_ = 0 for trait 2. With the current two-trait example, only three out of a total of four possible phenotypes can be achieved (Figure 1B).

Fitness is determined by a mapping function relating a phenotype to an environment (Equation (6)), Figure 1C. We assume, for simplicity, that for any specific environment there is a single optimal phenotype. This optimal phenotype is represented by a Boolean vector ***b*** whose element *b*_*i*_ is the optimal state of trait *i* in this environment. An individual’s absolute fitness is defined as the proportion of its traits that match the optimal phenotype.

Robustness in the model corresponds to the sensitivity of a phenotype to a single mutation in *υ*, manifested by the probability that a trait in *z* changes as a consequence of this mutation. The probability *P* that a trait *z*_*i*_ changes as a consequence of a mutation of *υ*_*i*_ is given in (Eq. 3). Genetic robustness in this formulation can be defined as 1 − *P*, or *n*(1 − *P*). Under the genotype-to-phenotype function, as *γ* increases over its range, so too does the number of traits it influences by a mutation (Figure 2). Therefore, *γ* is a robustness parameter in the model. For the purpose of allowing robustness to evolve without changing the phenotype (as it can with *γ*), a second robustness parameter, *α*, is introduced into the model (Eq. 4). We investigate robustness by varying either *γ* or *α*. Shown in Figures 2 and 3, the parameter *γ* has a narrower impact on robustness than the parameter *α*. For example, at the lowest level of robustness by *γ* (fixing *α* = 1), one mutation changes four traits out of 1000 on average, whereas under high values of *α* (fixing *γ* = 1), one mutation can change up to 250 traits out of 1000 (parameters given in Table 2).

**Table 2:** List of parameters

| parameters | definition | value(s) |
| --- | --- | --- |
| $c$ | probability of $t_{ij}$ not 0 | 0.5 |
| $a$ | proportion of 1s in $\mathbf{v}$ | 0.5 |
| $n$ | length of phenotype vector $\mathbf{z}$ | 1000 |
| $L$ | length of genotype vector $\mathbf{v}$ | 10000 |
| $\gamma$ | activation/repression coefficient of each gene on its target trait | 0.1, 1, 10 |
| $\alpha$ | a second parameter to scale robustness | 0.5, 1, 2, 2.5, 3, 3.5, 4, 8, 16, 64, 128, 256 |
| $\mu$ | mutation rate per gene | $10^{-6}$ |
| $N$ | population size | 10000 |
| $\mu_r$ | mutation rate on robust genotype | $10^{-5}$ |

**Figure 2:**
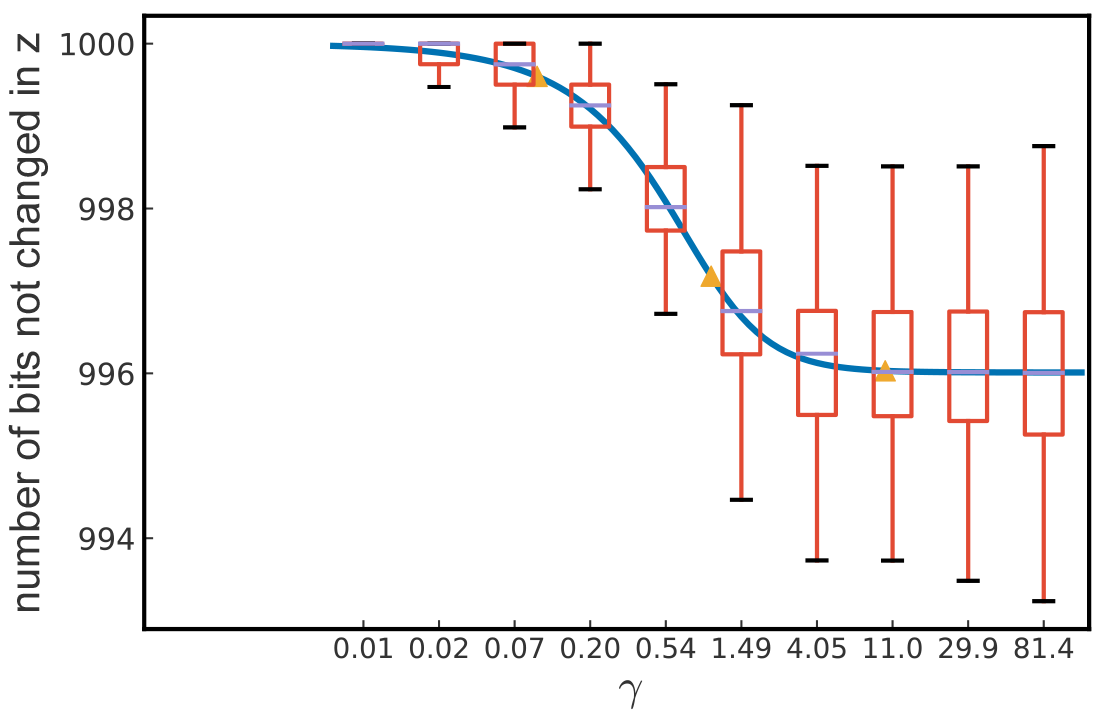
Control of robustness by *γ*. The number of traits that are not changed in phenotype *z* given a mutation in genotype vector ***υ*** with different *γ*, which is a measure of robustness. *c* = 0.5, *a* = 0.5, *L* = 10000, *n* = 1000. Boxplots are calculations from simulations and the solid line shows the analytic result (from 652 equation (3), showing *y* = *n*(1 - *P*)).For the simulations, we first simulate traits based on parameters. Then for each simulated trait, every bit in the genotype ***υ*** is flipped and the new phenotype is calculated, the number of bits in the new phenotype compared to the initial one that are kept the same is recorded. 1000 simulations are done for each parameter set. Yellow triangles mark three *γ* values: 0.1, 1, and 10, which are used for the following evolutionary simulations.

**Figure 3:**
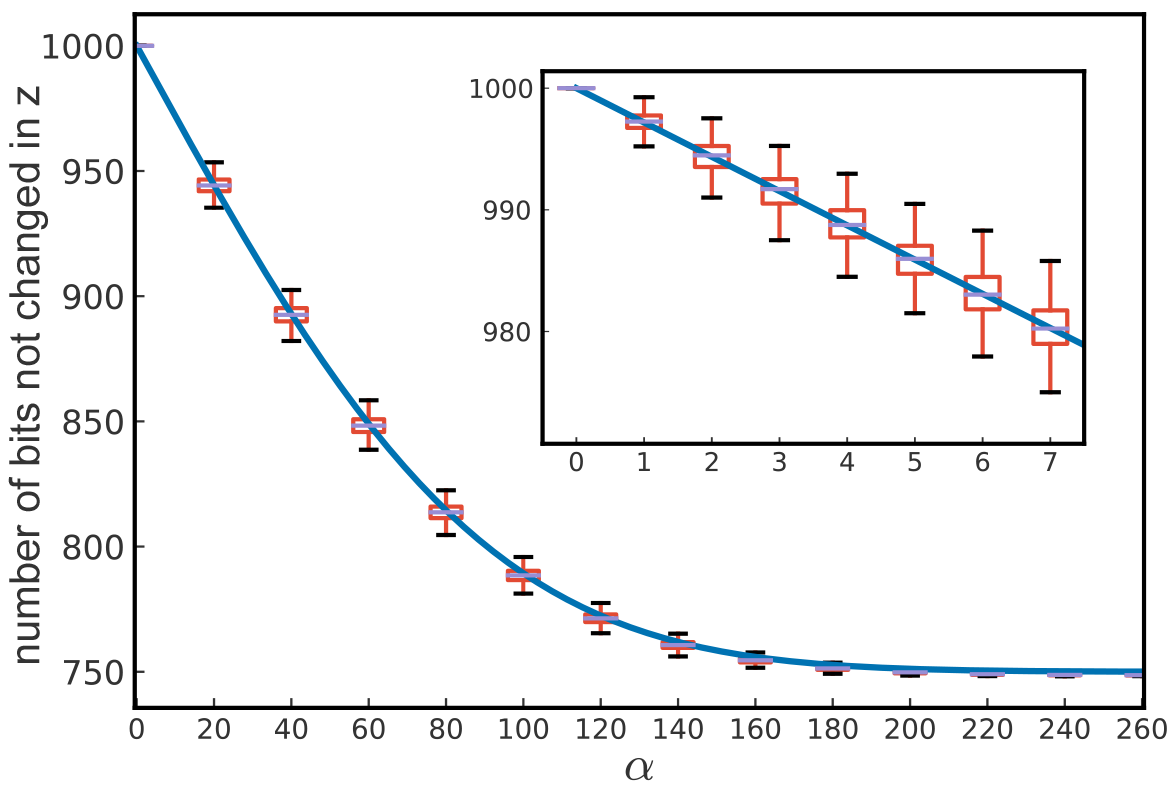
Control of robustness by *α*. The number of traits that are not changed in phenotype *z* given a mutation in genotype vector ***υ*** with different *α*. Boxplots are calculations from simulations while the solid line shows the analytic result (from equation (5), showing *y* = *n*(1 - *P*)). For the simulations, initial traits are simulated based on parameters in Table 2 and *γ* = 1. Then *α* is applied for each trait, and number of bits that are kept the same in phenotype when a bit change in ***υ*** is evaluated for each *α*. 1000 simulations are done for each parameter set. The nested small plot shows the relation to robustness for *α* <= 7, where an almost linear relationship is observed.

We evaluate evolution at different robustness levels by carrying out Wright-Fisher forward evolutionary simulation of a population by mutating the genotype vector, and monitoring population mean fitness as it adapts from its current phenotypic state. We consider two possible scenarios: (1) when the population is well adapted to its environment, which remains unchanged (E_opt_), and (2) when the population is challenged to adapt to novel environment, represented by a new optimal phenotype (E_new_). Values of parameters in the model used for this purpose are given in Table 2. We set the total number of genes *L* to be of the same order of magnitude (10, 000) as the number of genes in a typical eukaryote. Under this parameter regime, the individual probability of acquiring a new mutation each generation is 0.01; with the population size (*N* = 10, 000), 100 new mutations are introduced on average each generation into the population. Unless stated otherwise, these parameter values are used for all simulations (Table 2). The following two sections report results varying *γ* and *α*, corresponding to the evolution of populations across narrow (*γ*) and wide (*α*) ranges of robustness levels, respectively.

### A. Impact of *γ* on evolution of robustness

*γ* has a monotonic relationship with robustness—robustness decreases with increasing *γ*—but this relationship has upper and lower bounds (Figure 2). We selected three values of *γ*—0.1, 1, and 10, for subsequent use in simulations. The three values are well separated in the dynamic range of robustness to represent high, medium and low robustness, respectively (Figure 2). Note that the parameter *γ* has limited impact on the number of traits that change with the introduction of a single mutation (an average maximum of four traits).

#### a. Evolution in a constant, optimal environment (E_opt_)

We first examine evolution under an environment to which the population is initially fixed for the optimal genotype (E_opt_; Figure 4A). Models of robustness are often explored under this scenario, equivalent to stabilizing selection [Wagner et al., 1997, Eshel and Matessi, 1998, Wagner, 1996, Siegal and Bergman, 2002, Bullaughey, 2011].

**Figure 4:**
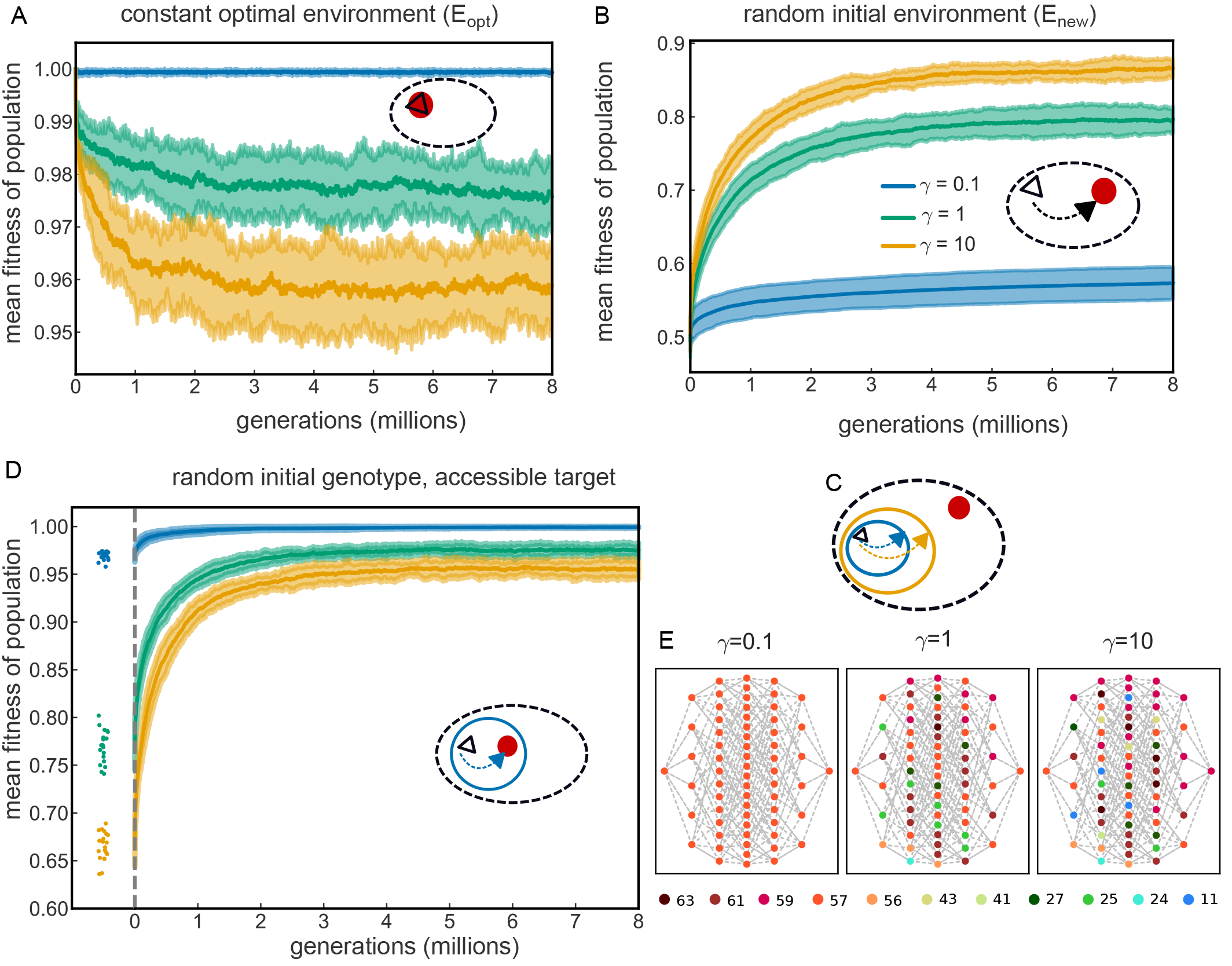
Different evolutionary trajectories with a fixed level of robustness. Schematic plots (oval-shaped graphs) in A-D indicate what kind of evolutionary regimes the simulations are in. Black dashed oval indicates the total phenotypic space. Black triangle represents the initial phenotype of the starting population. Red circle indicates the optimal phenotype specified for each environment. Solid circle with either blue or yellow color represents the total phenotypic space for *γ* = 0.1 and *γ* = 10 respectively. A. Mean fitness changes over time for populations with different robustness levels under optimal stable environment (shortened as E_opt_). 20 simulations were done for each *γ*. The thick solid lines show the mean of the 20 simulations at each generation. The upper and lower boundaries are 1.5 standard deviation on each side of the mean (same in B, D). *γ* = 0.1 in blue, *γ* = 1 in green and *γ* = 10 in yellow. Results from every 1000th generation are plotted due to large number of generation simulated (same in B, D). B. Mean fitness changes over time for populations with different robustness levels under a random initial environment (shortened as E_new_). C. Schematic plot illustrates high robustness population have a much smaller phenotypic space compared to low robustness ones. D. Mean fitness changes over time for populations initialized with a “attainable” random environment, where this new environment is initialized using a random genotype, and the same genotype to phenotype map to calculate the optimal environment b. The left of the plot shows the initial mean fitness of each population with a new “attainable” random environment. Each dots represents one simulation. E. One simulation showing phenotypic space for all genotype combinations for different robustness for *n* = 6, *L* = 6. Each circle represents a unique genotype, and each column indicates different total number of “1”s in the genotype vector (from left to right, zero “1”s to six “1”s). Each genotype is connected to its neighboring genotype that has 1 bit of difference in dashed lines. One set of randomly generated ***T*** and ***h*** was used for all *γ* values and for all combinations in genotype. For different *γ* values, the ***T*** was scaled to *γ****T***, according to the equation (1). The corresponding phenotype for each genotype was calculated. Each Boolean phenotype vector was then converted to an integer value, from which each unique value is assigned to a color, shown in the bottom.

Figure 4A plots mean population fitness across generations for three values of the robustness parameter *γ*. Populations are initially fixed for the optimal genotype and are allowed to evolve until a plateau in population mean fitness is observed. With high robustness (*γ* = 0.1), the population mean fitness remains close to the optimal value (*w̅* = 0.99929 ± 0.00025), while when robustness is reducd *γ* = 10), the population mean fitness equilibrates at a lower value (*w̅* = 0.95869 ± 0.00478). For populations close to their optimum, high robustness is selectively favored over low robustness, consistent with previous conclusions on robustness under stabilizing selection [Wagner et al., 1997, Eshel and Matessi, 1998, Wagner, 1996, Siegal and Bergman, 2002, Bullaughey, 2011]. Since the optimal phenotype is fixed in the initial population, mutations are deleterious initially (Figure S1C), and even at equilibrium the distribution of selective effects remains negatively skewed with a negative mean (Figure S1D). The reduction in population mean fitness is driven mostly by fixed mutations rather than segregating mutations (Figure S2). In fact, a greater proportion of mutations are purged from the population when robustness is low, resulting in lower mean population heterozygosity (Figure S1A). The loss of fitness when robustness is lower, we hypothesize, may be a consequence of genome-wide hitchhiking effects of selection against deleterious mutation under our model of complete linkage of loci [Charlesworth, 2013, Charlesworth et al., 1993, Haigh, 1978].

We also investigated evolution under E_opt_ for all combinations of values of *n* and *L* ∈ {100, 1000, 10000}, with all other parameters being the same (Figure S3). Within the robustness range determined by *γ*, there is an overall decline in population fitness with decreasing robustness, and the sensitivity of population fitness to robustness is greater with larger number of genes. One combination of parameter values (*n* = 10,000, *L* = 1000), however, produces an anomalous result in that this pattern reverses, with only high robustness having reduced fitness (*γ* = 0.1) (Figure S3F). This behavior may be a manifestation of Mullers Ratchet after the population exceeds a threshold number of deleterious mutations, initially hidden by robustness, at which point mean fitness declines precipitously.

#### b. Evolution following a sudden environmental shift (E_new_)

What will the relationship between robustness and evolution be when there is a sudden environmental shift? We first explore this question by randomly creating a new environment ***b***\* with its corresponding optimal phenotype (Materials and Methods for details), and then challenge the population, previously optimized for E_*opt*_, to evolve towards this new optimum. This leads to the expected initial mean fitness of population to be 0.5 according to our fitness definition (Figure 4B). We find that populations with low robustness (*γ* = 10) adapt more quickly than those with high robustness (*γ* = 0.1). Low robustness populations also reach substantially higher equilibrium mean fitness than high robustness populations (Figure 4B). Both of these observations—the initial rate and the final mean fitness of population—have monotonic relationships with robustness in the chosen parameter range of *γ* (Figure S4). The increase of population mean fitness is driven mostly by fixed mutations (Figure S2).

The aforementioned simulations were initialized with a genetically homogeneous population, whereas most populations will be genetically variable prior to an environmental shift. We therefore investigated whether initial genetic variability affects the ability of a population to evolve to a new optimum phenotype, noting that robust populations are more genetically variable than a less robust populations (Figure S1A). We initialized populations by subjecting them to 8×10^6^ generations of stabilizing selection under E_opt_ (Figure 4A) before applying a change in environment (a randomly chosen ***b***\*, E_new_) in the very next generation. We then allowed the populations to adapt to this new environment for another 8 × 10^6^ generations. The mean fitness change after the shift (Figure S1B) indicates little difference in the adaptive dynamics compared to populations starting fixed for a single genotype (Figure 4B vs. Figure S1B). Thus there is no initial advantage under this framework in highly robust populations to have higher genetic diversity in terms of their ability to adapt to a novel environment. This behavior occurs because the model specifies the same relation of phenotypic sensitivity to mutation at each robustness level. For populations with high robustness (*γ* = 0.1), even though they have greater initial genetic diversity, there is less phenotypic variation on which selection can act.

The distribution of selection coefficients, *s*, when adapting towards a new optimum, is initially close to symmetric under our model for high and medium robustness, with a positive mean for low robustness (Figure S1E). Moreover, low robustness populations have a larger variance in fitness, which accounts for their more rapid initial increase in fitness (as predicted by Fishers fundamental theorem of natural selection).

High robustness populations also reach a lower fitness plateau than low robustness popu lations. This is a surprising finding because the simulations show that highly robust populations do not evolve more slowly to eventually achieve the higher fitness peak attained by low robustness populations. One possibility is that high robustness populations become trapped with sub-optimal phenotypes because phenotypes with higher fitness cannot be realized due to constraints imposed by robustness. We hypothesized that high robustness results in a smaller phenotypic space than low robustness, and that when a random phenotype target is drawn from the universe of possible phenotypes, it will typically be outside of the realizable phenotypic range of the population. Since low robustness explores a larger phenotypic space, it can evolve to a phenotype closer to the random phenotypic target (Figure 4C).

To test this hypothesis, we set up the initial “random” optimal environment ***b***\*\* such that its corresponding optimal phenotype is reachable within each population’s genotypic search space (Figure 4D). Under this scheme, populations achieved similar mean fitness for each robustness level as when they started with the optimal phenotype (Figure 4A, D), indicating that when the phenotypic target is achievable within their phenotypic space, populations under all three robustness levels evolve to their new optimum and reach a mutation-selection balance.

High robustness constrains mutant individuals to have a phenotype more similar to their parental genotypes than low robustness (Figure 4D, initial fitness), which further supports our hypothesis. Since it is not practical to enumerate the complete phenotypic space for *L* = 10000 genotypes, we explored this point by enumerating all possible genotypes for six genes (*L* = 6), and calculating phenotypes produced by these genotypes with the same genotype to phenotype map. This confirms, at least in low dimension, that high robustness produces a much smaller phenotypic space than low robustness (one simulation example shown in Figure 4E).

### B. Impact of *α* on evolution of robustness

The phenotypic effects of mutations regulated by *γ* are restricted to a relative small range constrained by the model (Figure 2). Varying alpha allows us to explore a wider range of robustness (Figure 3). As before, we investigate alpha for populations under E_opt_ and E_new_ (with *γ* set to 1).

Rather than the monotonic relationship of robustness to fitness found when varying *γ* (Figure 4A, B), we observe a bifurcation in final mean fitness at intermediate *α*, and an inversion in the relationship between robustness and evolution when robustness decreases further (larger *α*). Simulations with different *α* were run for twice the number of generations as in previous simulations to assure that asymptotic population fitness was achieved (1.6×10^7^ total). Typical trajectories are summarized in Figure 5A and 5B (see Figure S5, S6 for all *α* values), showing two representative simulations for four alpha values (*α* = 1, 2, 3, 8) either under E_opt_ or E_new_. Mean fitness at the end of a simulation is shown for each *α* value in Figure 5C and 5D. When *α* is no larger than 2.5, the behavior of populations recapitulates the patterns described in the previous section for robustness regulated by *γ* (grey shaded area show the corresponding robustness range explored by *γ* in Figure 5C and 5D). However, as robustness decreases further (3 <= *α* <= 4), population fitness trajectories bifurcate. For E_opt_, populations either remain close to the optimal fitness value, or drop to a lower fitness value (Figure 5A, C). When challenged to adapt to E_new_, populations either reach a higher fitness value or are trapped with sub-optimal phenotypes (Figure 5B, D). When robustness decreases even further (*α* >= 8), populations remain close to the optimal fitness in E_opt_, but are unable to reach high fitness when challenged with E_new_ (Figure 5C, D). Interestingly, the initial adaptive behavior of populations with respect to *α* differs from the final adaptive state of the populations in E_new_. Specifically, we find that low robustness populations, though they eventually adapt poorly to a novel environment, initially adapt more quickly than high robustness populations (Figure 5D, E).

**Figure 5:**
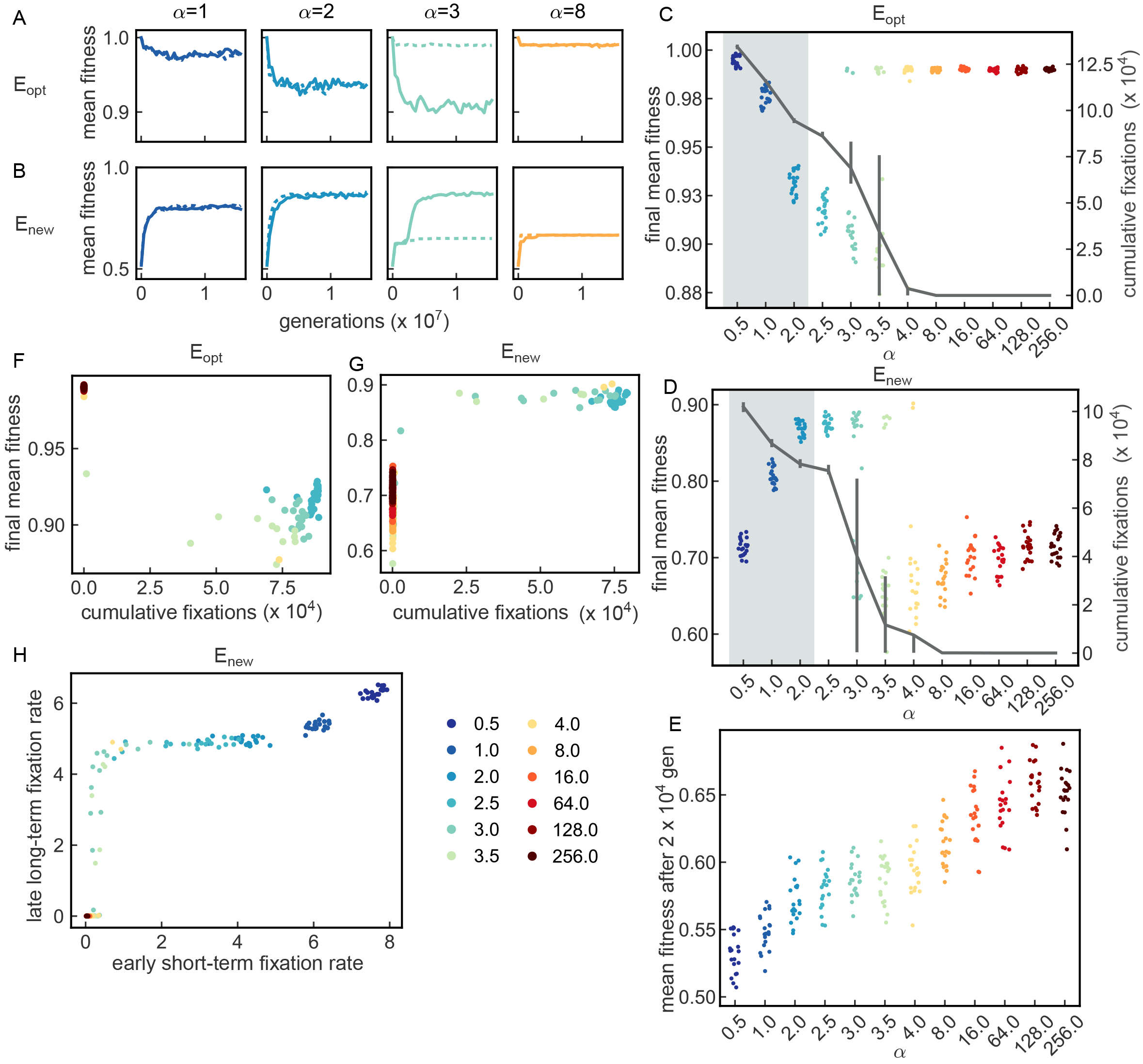
Populations showing non-linear relationship with larger range of robust-ness under constant optimal or random initial environment. A. Two representative simulations showing mean fitness of population changes over time for *α* values 1, 2, 3 and 8 under E_opt_. One simulation is in solid line and the other is in dashed line to distinguish, over 1.6 ×10^7^ generations. (Results from every 400,000th generation are plotted due to large number of generation simulated. Same in B.) B. Two representative simulations showing mean fitness of population changes over time for *α* values 1, 2, 3 and 8 under E_new_. C. Each dot represents the final population mean fitness at the generation of 1.6×10^7^ from one simulation under E_opt_. Different colors represent different alpha values. Jittering is used to display points with the same *α* values. Note this color scheme for different *α* values are used consistently also in Figure 6 and Figure 7. The grey line shows the mean number of fixations at the end of the simulations with one particular *α* values. The error bars are showing 15% and 85% of the final fixation numbers. The light grey rectangle-shaded area shows the similar parameter range of robustness as *γ*. D. Same as C under E_new_. E. Mean fitness of each population after 2 ×10^4^ generations under E_new_. F. The relationship of the cumulative number of fixations to the final mean fitness of population under E_opt_. Only for *α* values from 2.5 to 256. G. Similar as E under E_new_. H. Fixation rate is calculated as the average number of fixations per 1000 generations given a period of time. Early short-term fixation rate is the average fixation rate for the initial 10^6^ generations, and the late long-term fixation rate is the average fixation rate from 1000001 generation to 1.6 × 10^7^ generation.

We sought to find the cause of the bifurcation at intermediate *α* and the inverted behavior when robustness decreases further. We first examined the total number of fixations for each *α* value at the end of the simulations (Figure 5C, D). We find a similar pattern for the cumulative number of fixations for E_opt_ and E_new_ simulations—total fixations decrease with decreasing robustness. We also observed an increased variance in the number of fixations for *α* values between 3 and 4, which correlates with the bifurcation behavior in the final mean fitness of each population. We also compared the cumulative number of fixations against final population mean fitness under both E_opt_ and E_new_ for *α* >= 2.5 (Figure 5F, G). We observe two non-overlapping clusters of population final mean fitness, one high and one low, separated by fixation number. For E_opt_, populations that achieve high fitness fix no mutations (Figure S8), while the ones that fix many mutations have a lower fitness. For E_new_, populations that fix few mutations remain at a low fitness while ones with many fixations achieve a higher fitness. Note for E_new_ at *α* = 8, there are fixations in the first 2 × 10^6^ generations but populations nevertheless can remain trapped at low fitness (Figure S9).

In Figure 5H, the fixation rate (the average number of substitutions per 1000 generations) in the first 10^6^ generations is plotted against the next 15 × 10^6^ generations (early vs. late, respectively). When *α* <= 2, the fixation rate is high and differs little between the two time intervals. When *α* >= 8, both early and late fixation rates are close to 0. In contrast, for intermediate alpha values (2.5 <= *α* <= 4) the substitution rate is low early and late, or it can be low early but jump to a higher value in the longer second interval. This latter situation represents evolutionary trajectories leading to high fitness (but often not until the second time interval). Whether or not populations can gain a positive long-term fixation rate determines the bifurcation behavior.

#### a. A co-evolved locus that controls robustness

To gain a better insight of how robustness could evolve indirectly when only the phenotype is under direct selection, we introduce a mutable locus, ***r***, controlling *α* and completely linked to the genotype ***υ***. The reason we chose to change *α* rather than *γ* is because it has the desirable property of only affecting sensitivity of mutations on phenotypes while keeping the current phenotype the same. We chose to have alleles in ***r*** encoding eight levels of robustness, ranging from 0.5 to 8 (the first eight alpha values in Figure 5C), using three bits to represent the eight states (Table S1). A mutation in the robustness locus alters a single bit, as is the case for the rest of the genotype. The mutation rate of the robustness locus is set to *μ*_*r*_ = 10^−5^. Fitness is calculated as previously described (Eqs 1, 4, 6). The population is initialized with equal frequencies of the eight robustness alleles. The population then evolves under the same two environmental scenarios as before (E_opt_ and E_new_).

Our initial simulations were carried out with the robustness locus completely linked to genotype ***υ***. Under E_opt_, the most robust genotype goes to fixation in every simulation we examined (Figure 6A, B). This behavior is consistent with mean population fitness declining with decreasing robustness when *α* <= 2.5 (Figure 5A, C). We note, however, that high mean population fitness is achieved on either side of the alpha bifurcation, with the least robust population (*α* = 8) actually having the highest fitness. Yet, these simulations invariably fix instead for the most robust allele. When examining the population dynamics with constant alpha in the initial 200 generations in E_opt_, populations with the higher robustness do have less deleterious mutations (Figure S10).

**Figure 6:**
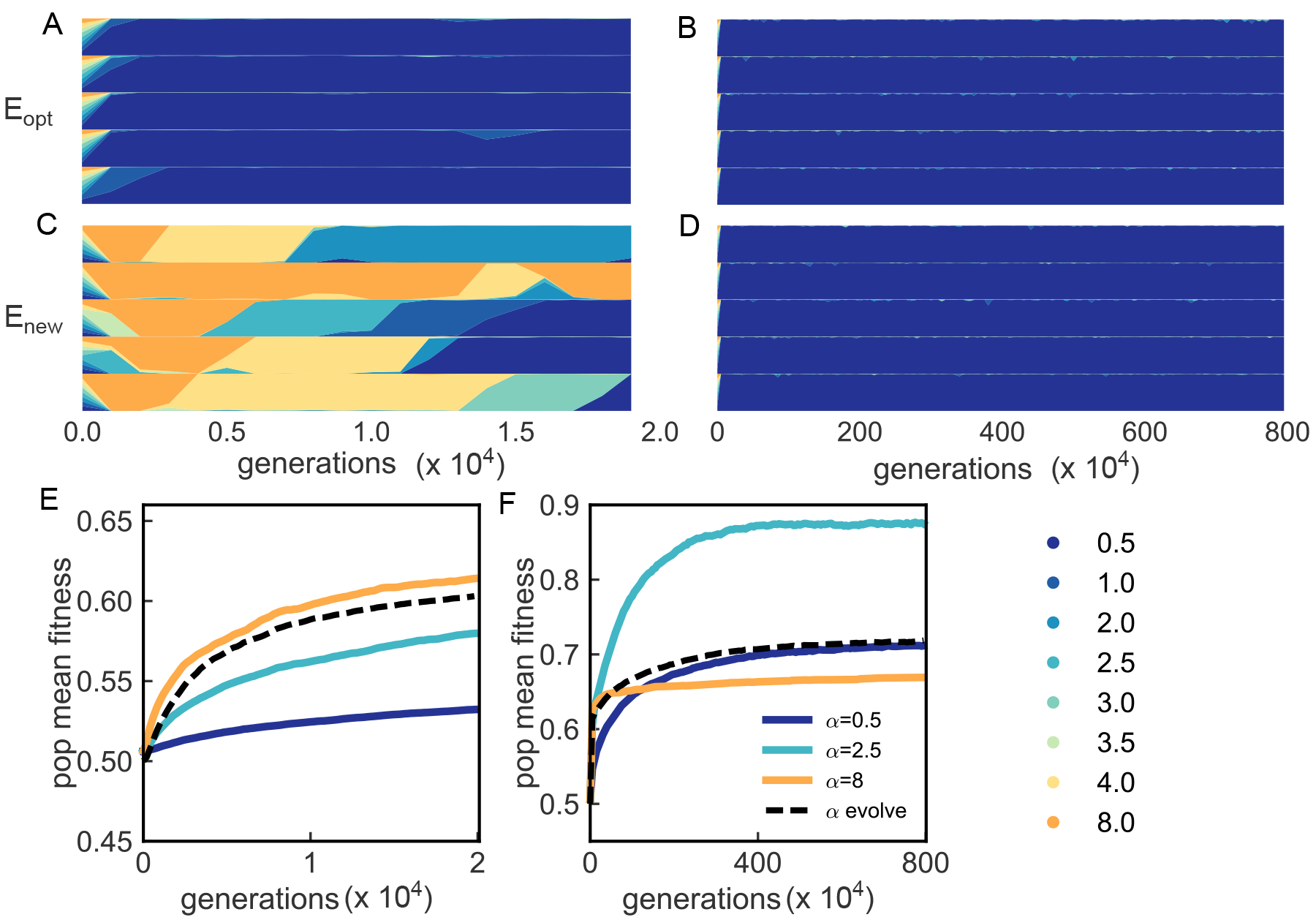
Population dynamics with evolving robustness. Simulations of populations with evolving robustness. Only 5 representative simulations are shown for A-D. For A and C, robustness genotype frequency from every 1000th generation are plotted. For B and D, robustness genotype frequency from every 50,000th generation are plotted. A, B show the robustness genotype frequency changes over time under E_opt_, with A showing the change over the first 2 × 10^4^ generations, and B for 8 × 10^6^ generations. C, D show the robustness genotype frequency changes over time under E_new_, with C showing the change over the first 2 × 10^4^ generations, and D for 8 × 10^6^ generations. E-F Mean fitness change comparison for populations with fixed level of robustness and ones with evolving robustness under E_new_. Solid line shows the mean value of the mean fitness of 20 simulations. E shows the comparison for the the first 2 × 10^4^ generations and every 200th generation is plotted. F shows the comparison for 8 × 10^6^ generations and every 50,000th generation is plotted.

For populations adapting to a novel environment (E_new_), the lowest robustness allele is selectively favored initially (Figure 6C), consistent with initial population dynamics with constant robustness under this environment (Figure 5E). In Figure 6E, where for the first 2 × 10^4^ generations, the adaptive dynamics of the populations with evolving robustness is similar to the one with fixed *α* = 8. However, as the adaptive process continues, the highest robustness genotype (*α* = 0.5) takes over the population permanently. Population mean fitness then proceeds to increase along the trajectory given for fixed *α* = 0.5, to a stable mean population fitness of ≈ 0.7 (Figure 6F). Thereafter, it does not improve any further. Surprisingly, this is not nearly the highest fitness the population could have achieved. For example, with fixed *α* = 2.5 mean fitness rises to nearly *w̅* = 0.9. (Figure 6F). The high robustness genotype fixed early despite the fact that it prevents the population from evolving to a higher fitness.

We hypothesized that this behavior could depend on the specific parameterization or assumptions of the model. We investigated these possibilities by varying the mutation rate of the robustness locus ***r***, and also allowing the robustness locus to recombine with ***υ***. We found that decreasing or increasing mutation rates at the robustness locus increases the equilibrium mean fitness (Figure 7A). A lower mutation rate prolongs the time to elimination of low robustness alleles, allowing greater time to select for a more highly adapted phenotype (Figure 7C). A higher mutation rate increases the number of segregating robustness alleles (Figure 7D). Recombination between the robustness locus and ***υ*** also increases the final mean fitness of the population (Figure 7B). Indeed, the highest mean fitness is reached when at the maximum recombination rate (*r* = 0.3) between ***r*** and ***υ*** in the simulation (Figure 7E). These results show that the dynamics of adaptation when robustness can also simultaneously evolve is sensitive to the genetic parameters (*i.e*., mutation and recombination rates) of the system.

**Figure 7:**
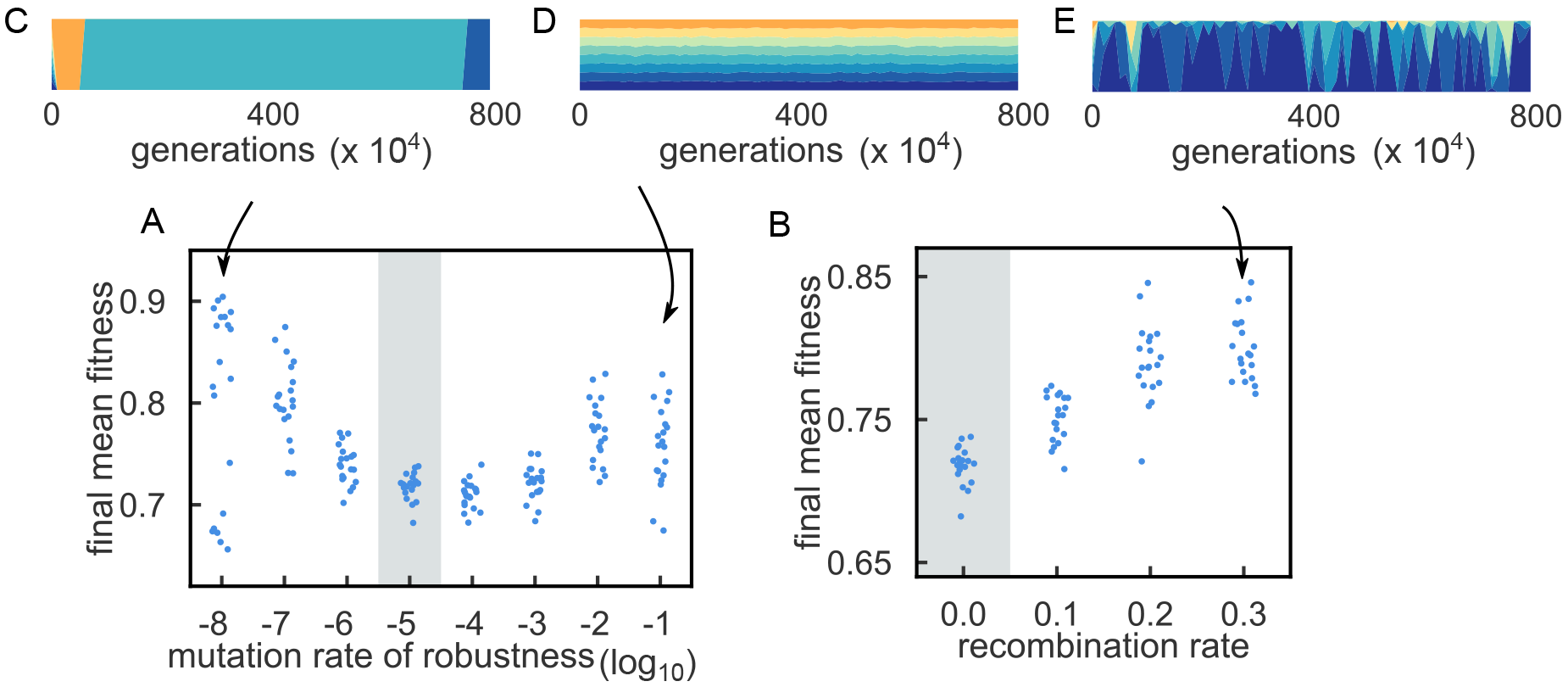
Mutation and recombination rate of robustness locus affects the adap-tation process. Mutation rate of robustness locus and rate of recombination between robustness locus and genotype ***υ*** are explored under E_new_. A. Final mean fitness of population after 8 × 10^6^ generations with different mutation rate of the robustness locus. Each dot represent one simulation result. Jittering is applied for simulations with the same mutation rate. B. Final mean fitness of population after 8 × 10^6^ generations with different recombination rate between the robustness locus and genotype v. Jittering is applied. The recombination rate ***r*** is the probability that a recombination event occur between randomly chosen two haploid. C. One example simulation of robustness genotype frequency changes over time, with mutation rate of robustness genotype to be 10^−8^. D. One example simulation of robustness genotype frequency changes over time, with mutation rate of robustness genotype to be 0.1. E. One example simulation of robustness genotype frequency changes over time, with recombination rate between robustness locus and the regular genotype to be 0.3 per individual per generation.

## Discussion

Our model and simulations explored the relationship between robustness and evolvability. Two models were investigated, one in which robustness is set at fixed values of *γ* and *α* and evolutionary dynamics are compared across a range of robustness levels (Figure 4, 5), and the other in which robustness is controlled by a mutable locus and both phenotype and robustness are allowed to evolve simultaneously (Figure 6). Two environmental regimes were investigated, selection around a phenotypic optimum (E_opt_) and selection for a novel phenotype (E_new_), representing situations analogous to long-term environmental stasis or alternatively rapid environmental change (Punctuated equilibrium [Gould and Eldredge, 1977, Eldredge and Gould, 1972]).

Robustness, defined as the probability that a trait does not change with a mutation, is constrained at its low end by the mathematical formulation of the genotype to phenotype map (Equation (3), (5), Figure 2, 3). For large *α*, a maximum of approximately one quarter of the traits can change with a mutation (Figure 3); For large *γ*, this value is about 70-fold lower (Figure 2). The two robustness parameters, therefore, extend over distinctly different ranges of robustness. With the advantage of the explicit representation of genotype to phenotype in our model, we could associate the level of robustness to its biological meaning rather than an abstract parameter.

Robustness is generally evaluated by its impact on evolvability. Different conclusions can be reached, however, depending on how evolvability is defined. In our simulations, the robustness parameter can be seen to affect both the initial rate of adaptation and the final population mean fitness, suggesting two different definitions for evolvability. Consider the robustness range regulated by *α*. If considering the initial rate of adaptation, evolvability increases monotonically as robustness decreases (Figure 5E). If considering the final population mean fitness instead, its relationship to robustness is more complex. When *α* is relatively small, evolvability increases as robustness decreases, bifurcates at intermediate *α*, and collapses to low values at even larger *α*. Our simulations highlight the complexity of the adaptive process with respect to robustness, indicating the need to be precise in defining evolvability, and the value of studying its temporal dynamics.

Our study shows that conclusions about the relationship between robustness and evolvability also depend on the realizable range of robustness. In biological systems, genetic components have different limits in regulating robustness. For example, mutations changing the *cis*-regulatory elements of a gene usually have a smaller phenotypic effect than those that change protein coding regions. Mutations in certain genes, such as *Tabby* or *scute*, affect phenotypic variation in specific traits, whereas Hsp90 mutants tend to affect many traits simultaneously.

One frequently posed question regarding how canalized traits evolve concerns whether a phase of decanalization (a decrease in robustness in our model) is required. Our results generally affirm this view and provide insights into how different levels of robustness can be favored depending on the adaptive scenario. With the parameter *γ*, under E_opt_, high robustness is selectively favored (Figure 4A), indicating that canalization (high robustness) should be prevalent under conditions such as stabilizing selection for an optimal phenotype. E_new_, in contrast, imposes a sudden reduction in fitness and forces the current population to adapt to a new optimum. We found that this scenario favors low robustness (Figure 4B). In the absence of change in the robustness level, canalized traits may have limited opportunity to adapt. We found that populations with high robustness, although they accumulate genetic variation under long-term stabilizing selection, still cannot achieve high mean fitness without a decrease in robustness (Figure S1). Simulations in which robustness is encoded as a mutable locus allowed both phenotype and robustness to evolve simultaneously over a larger robustness range set by *α*. These simulations also find an initial selective advantage of low robustness and a long-term selective advantage of high robustness (Figure 6A-D).

Experimental evidence is consistent with the conditions in our model that favor low or high robustness. First, low robustness at the onset of environmental change has been observed in diverse biological systems ([Bjedov et al., 2003, Galhardo et al., 2007, Radman, 1974, Echols, 1981]), suggesting that low robustness is preferred in the initial stage of adaptation. *E. coli* cells under certain stress conditions increase their mutation rate by activating the SOS pathway, changing to error-prone polymerase [Bjedov et al., 2003, Galhardo et al., 2007], creating DNA-damage, and increasing recombination [Radman, 1974, Echols, 1981]; Stress-induced mutagenesis can be interpreted as an adaptation to decrease robustness in response to severe environmental change.

Second, studies of natural populations find evidence favoring high robustness when the population is near or approaching its optimum. Hsp90 buffers standing genetic variation in yeast populations, but displays the opposite effect for de novo mutations in mutation accumulation lines [Geiler-Samerotte et al., 2016], indicating that this chaperone protein enhances robustness in stable populations. Similarly, study on the yeast TDH3 promoter shows polymorphisms have less expression noise than random mutations, which indicates that natural selection maintain mutations with high robustness [Metzger et al., 2015].

The simulations revealed limits on the ability of selection to evolve to a new optimal phenotype, especially under high robustness (Figure 4B). We attribute this constraint on evolvability to the much smaller set of possible phenotypes achievable in the model under high robustness (Figure 4E). This aspect of evolvability is generally not considered in population genetic models, but, we believe, is a biologically realistic property of developmentally regulated phenotypes. Loci such as Hsp90 can regulate the size of the universe of realizable phenotypes by changing from high robustness to low robustness by mutations, indicating that robustness may itself be an evolvable trait.

An unexpected result is the bifurcation behavior for *α* values in the range of 3 <= *α* <= 4. In this range of *α*, a single mutation will cause approximately 8 to 12 traits to change states (Figure 3, inset). Populations having the same parameter set in this range can take one of two different fates. In E_opt_, population final mean fitness declines as *α* increases from 0.5 to 2.5, indicating that weak but increasingly deleterious mutations are being fixed in the populations by genetic drift. To the right of the bifurcation zone (*α* > 4), final population mean fitness remains high, close to 1, and the number of fixations decreases to near-zero. At large *α*, mutations become universally deleterious and are selectively eliminated because many traits are simultaneously changed. Mutation-selection balance, in this case, keeps populations near their optima.

The same logic can be applied to E_new_, where in this case, except the very initial phase of adaptation, mutations under large *α* are deleterious, and the population fails to adapt to a novel optimum, and again the number of fixations is near-zero. At small *α* values (0.5 <= *α* <= 2.5), however, increasing *α* to cause a greater number of traits to change allows populations to adapt better to the novel optimum. In the bifurcation zone, either fate is possible, suggesting that adaptation is contingent on the existence of critical beneficial mutation(s) that then allow the population to continue evolving adaptively over a longer period of time.

For simulations in which robustness is encoded as a mutable locus allowing both phenotype and robustness to evolve simultaneously, we discovered that high robustness alleles could be selectively favored relatively early on, hindering the population from achieving a higher mean fitness (compared to simulations with a constant but slightly lower robustness level). This is another way in which high robustness might constrain adaptation. Upon further investigation, this constraint was found to be dependent on the lack of recombination between and these *cis*-regulatory sites, and the transcriptional unit whose expression they regulate, are tightly linked genetically. Increasing the recombination distance between *cis*-regulatory modules and the transcriptional unit should, therefore, be selectively favored. There may be a selective advantage, therefore, for spreading of cis-regulatory modules across a larger physical genomic landscape.

Our model employs an unconventional definition of phenotype that we believe can have broad utility in modeling the evolution of a complex genetic trait (with or without consideration of robustness). Phenotype, in the model, is simply a fixed-length vector of discrete traits, each trait having output states that are determined by the additive or subtractive activities of genes acting as *trans*-regulators of trait expression and a threshold. This genetic model of traits mimics developmentally regulated terminal cell type differentiation controlled by the expression state of regulatory genes. Other examples include the gene regulatory network for the determinationn sensory organ fate [Cassidy et al., 2013], epidermal cell fate [Chanut-Delalande et al., 2006], abdominal pigementation in fruit flies [Rogers et al., 2014], wing-spot patterning in butterflies [Gompel et al., 2005, Werner et al., 2010], and cell fate determination from stem cells [Huang et al., 2015, Wagers et al., 2002].

The representation of phenotype as a vector of distinct traits with defined genetic underpinnings has utility from an evolutionary perspective as well. Phenotype, in our model is the complete set of traits that are under selection. Similarly, selection in nature may act on a plethora of seemingly unrelated traits. For example, to increase insect flight duration or maneuverability, many distinct traits may be simultaneously subject to selection, including wing morphology, body size, flight musculature, sensory perception, energy metabolism, etc. Finally, the model incorporates both genetic pleiotropy (*c* = 0.5) and consequent correlations among traits.

Our model can be extended to consider more complex biological scenarios. The genotype vector maps directly onto a phenotype vector in this work. A more realistic scenario for multi-cellular organisms would be to model a hierarchy of gene regulatory networks [Erwin and Davidson, 2009]. Our model can be adapted to have two-layers translating genotype to phenotype, with the first layer incorporating a function for mapping genotype onto gene expression only. A second layer could then retain the function in our current model for map ping gene expression state onto the terminal phenotype. The current model can be adapted, moreover, to incorporate phenotypic plasticity by taking in environmental signals. In this case, an additional layer could also be added to incorporate post-translational regulation of phenotype in response to environmental signals. Note that in our current model, the same set of phenotypes are under selection before and following a shift in the optimum phenotype. It would be interesting to explore an alternative scenario, whereby selection would act on previously hidden mutable dimensions of a phenotype before an environmental shift.

We held the threshold, ***h***, constant throughout the simulations, though like *γ* and *α*, it can also effect robustness. In experimental gene-regulatory networks, it is possible to manipulate thresholds in networks with miRNA. Specifically, mir-9a in *Drosophila* has been shown to regulate differentiation and robustness of bristle numbers [Cassidy et al., 2013, Posadas and Carthew, 2014]. It may thus be possible to test predictions of the model in experimental systems by manipulating thresholds.

Our model displays a rich dynamical behavior for investigating the conditions under which robustness is favored or disfavored evolutionarily. The model treats phenotype explicitly, and permits robustness to be explicitly controlled. The incorporation of these ideas into classical population genetic models, as we attempt to do here, is, we believe, an important direction to take in the future.

## Acknowledgment

Thanks Kevin Thornton and Jaleal Sanjak for help and explanation with fwdpp code. Thanks Si Tang and Alexandre Ramos for useful discussions on the model. Thanks Andrew Bergen for useful discussions with connection to population genetics.

## References

L. W. Ancel. A quantitative model of the simpson–baldwin effect. Journal of theoretical biology, 196(2):197–209, 1999.

N. Barkai and S. Leibler. Robustness in simple biochemical networks. Nature, 387:913–917, 1997.

A. Barrière, K. L. Gordon, and I. Ruvinsky. Distinct functional constraints partition sequence conservation in a cis-regulatory element. PLoS Genetics, 7:e1002095, 2011.

I. Bjedov, O. Tenaillon, B. Gerard, V. Souza, E. Denamur, M. Radman, F. Taddei, and I. Matic. Stress-induced mutagenesis in bacteria. Science, 300(5624):1404–1409, 2003.

J. D. Bloom, S. T. Labthavikul, C. R. Otey, and F. H. Arnold. Protein stability promotes evolvability. Proceedings of the National Academy of Sciences, 103(15):5869–5874, 2006.

K. Bullaughey. Changes in selective effects over time facilitate turnover of enhancer sequences. Genetics, 187:567–582, 2011.

J. J. Cassidy, A. R. Jha, D. M. Posadas, R. Giri, K. J. Venken, J. Ji, H. Jiang, H. J. Bellen, K. P. White, and R. W. Carthew. mir-9a minimizes the phenotypic impact of genomic diversity by buffering a transcription factor. Cell, 155(7):1556–1567, 2013.

H. Chanut-Delalande, I. Fernandes, F. Roch, F. Payre, and S. Plaza. Shavenbaby couples patterning to epidermal cell shape control. PLoS biology, 4(9):e290, 2006.

B. Charlesworth. Background selection 20 years on the wilhelmine e. key 2012 invitational lecture. Journal of Heredity, 104(2):161–171, 2013.

B. Charlesworth, M.T. Morgan, and D. Charlesworth. The effect of deleterious mutations on neutral molecular variation. Genetics, 134(4):1289–1303, 1993.

A. Crombach and P. Hogeweg. Evolution of evolvability in gene regulatory networks. PLoS Computational Biology, 4(7):e1000112, 2008.

J. F. Crow. 2. the concept of genetic load: a reply. American journal of human genetics, 15(3): 310, 1963.

T. D. Cuypers, J. P. Rutten, and P. Hogeweg. Evolution of evolvability and phenotypic plasticity in virtual cells. BMC evolutionary biology, 17(1):60, 2017.

J. A. Draghi, T. L. Parsons, G. P. Wagner, and J. B. Plotkin. Mutational robustness can facilitate adaptation. Nature, 463(7279):353–355, 2010.

R. B. Dun and A. S. Fraser. Selection for an invariant charactervibrissa numberin the house mouse. Nature, 181(4614):1018–1019, 1958.

H. Echols. Sos functions, cancer and inducible evolution. Cell, 25(1):1–2, 1981.

I. M. Ehrenreich and D. W. Pfennig. Genetic assimilation: a review of its potential proximate causes and evolutionary consequences. Annals of botany, 117(5):769–779, 2015.

N. Eldredge and S. J. Gould. Punctuated equilibria: an alternative to phyletic gradualism. In T. J.M. Schopf, editor, Models in Paleobiology, pages 82–115. Freeman, Cooper, San Francisco, 1972.

D. H. Erwin and E. H. Davidson. The evolution of hierarchical gene regulatory networks. Nature Reviews Genetics, 10(2):141–148, 2009.

I. Eshel and C. Matessi. Canalization, genetic assimilation and preadaptation: a quantitative genetic model. Genetics, 149(4):2119–2133, 1998.

M. A. Félix and M. Barkoulas. Pervasive robustness in biological systems. Nature Reviews Genetics, 16(8):483–496, 2015.

R. A. Fisher. The genetical theory of natural selection: a complete variorum edition. Oxford University Press, 1930.

N. Frankel, G. K. Davis, D. Vargas, S. Wang, F. Payre, and D. L. Stern. Phenotypic robustness conferred by apparently redundant transcriptional enhancers. Nature, 466:490–493, 2010.

R. S. Galhardo, P. J. Hastings, and S. M. Rosenberg. Mutation as a stress response and the regulation of evolvability. Critical Reviews in Biochemistry and Molecular Biology, 42(5):399–435, 2007.

K. A. Geiler-Samerotte, Y. O. Zhu, B. E. Goulet, D. W. Hall, and M. L. Siegal. Selection transforms the landscape of genetic variation interacting with hsp90. PLoS Biology, 14(10):e2000465, 2016.

G. Gibson. Decanalization and the origin of complex disease. Nature Reviews Genetics, 10(2): 134–140, 2009.

G. Gibson and I. Dworkin. Uncovering cryptic genetic variation. Nature Reviews Genetics, 5(9): 681–690, 2004.

G. Gibson and D. S. Hogness. Effect of polymorphism in the drosophila regulatory gene ultrabithorax on homeotic stability. Science, 271(5246):200, 1996.

N. Gompel, B. Prud’homme, P. J. Wittkopp, V. A. Kassner, and S. B. Carroll. Chance caught on the wing: cis-regulatory evolution and the origin of pigment patterns in drosophila. Nature, 433 (7025):481–487, 2005.

S. J. Gould and N. Eldredge. Punctuated equilibria: the tempo and mode of evolution reconsidered. Paleobiology, 3(02):115–151, 1977.

D. Grigoriev, J. Reinitz, S. Vakulenko, and A. Weber. Punctuated evolution and robustness in morphogenesis. Biosystems, 123:106–113, 2014. doi: doi:10.1016/j.biosystems.2014.06.013. PMCID in process; PMID: 24996115.

Gaël Guennebaud, Benoît Jacob, et al. Eigen v3. http://eigen.tuxfamily.org, 2010.

J. Haigh. The accumulation of deleterious genes in a populationmuller’s ratchet. Theoretical Population Biology, 14(2):251–267, 1978.

E. J. Hayden, E. Ferrada, and A. Wagner. Cryptic genetic variation promotes rapid evolutionary adaptation in an rna enzyme. Nature, 474(7349):92–95, 2011.

B. Huang, G. Li, and X. H. Jiang. Fate determination in mesenchymal stem cells: a perspective from histone-modifying enzymes. Stem cell research & therapy, 6(1):35, 2015.

S. Huang. The molecular and mathematical basis of waddington’s epigenetic landscape: A framework for post-darwinian biology? BioEssays, 34(2):149–157, 2012.

David Kirk. Saturation curve analysis and quality control. Shot Peener, 20(3):24, 2006.

S. F. Levy and M. L. Siegal. Network hubs buffer environmental variation in Saccharomyces cerevisiae. PLoS Biology., 6:e264, 2008. doi:10.1371/journal.pbio.0060264.

Manu, S. Surkova, A. V. Spirov, V. Gursky, H. Janssens, A. Kim, O. Radulescu, C. E. VanarioAlonso, D. H. Sharp, M. Samsonova, and J. Reinitz. Canalization of gene expression in the Drosophila blastoderm by gap gene cross regulation. PLoS Biology, 7:e1000049, 2009a. doi:10.371/journal.pbio.1000049 PMCID: PMC2653557.

Manu, S. Surkova, A. V. Spirov, V. Gursky, H. Janssens, A. Kim, O. Radulescu, C. E. VanarioAlonso, D. H. Sharp, M. Samsonova, and J. Reinitz. Canalization of gene expression and domain shifts in the Drosophila blastoderm by dynamical attractors. PLoS Computational Biology, 5: e1000303, 2009b. doi:10.1371/journal.pcbi.1000303 PMCID: PMC2646127.

Manu, Michael Z. Ludwig, and Martin Kreitman. Sex-specific pattern formation during early Drosophila development. Genetics, 194:163–173, 2013.

J. Masel. Evolutionary capacitance may be favored by natural selection. Genetics, 170(3):1359–1371, 2005.

J. Masel and M. L. Siegal. Robustness: mechanisms and consequences. Trends in genetics, 25(9): 395–403, 2009.

C. Mayer and T. F. Hansen. Evolvability and robustness: A paradox restored. Journal of Theoretical Biology, 430:78–85, 2017.

C. D. Meiklejohn and D. L. Hartl. A single mode of canalization. Trends in Ecology and Evolution, 17(10):468–473, 2002.

B. P. Metzger, D. C. Yuan, J. D. Gruber, F. Duveau, and P. J. Wittkopp. Selection on noise constrains variation in a eukaryotic promoter. Nature, 521(7552):344–347, 2015.

N. Moris, C. Pina, and A. M. Arias. Transition states and cell fate decisions in epigenetic landscapes. Nature Reviews Genetics, 17(11):693–703, 2016.

A. B. Paaby and M. V. Rockman. Cryptic genetic variation: evolution’s hidden substrate. Nature Reviews Genetics, 15(4):247–258, 2014.

K. Palacio-López, B. Beckage, S. Scheiner, and J. Molofsky. The ubiquity of phenotypic plasticity in plants: a synthesis. Ecology and evolution, 5(16):3389–3400, 2015.

M. Perry, A. N. Boettiger, J. P. Bothma, and M. Levine. Shadow enhancers foster robustness of Drosophila gastrulation. Current Biology, 20:1562–1567, 2010.

D. M. Posadas and R. W. Carthew. Micrornas and their roles in developmental canalization. Current Opinion in Genetics & Development, 27:1–6, 2014.

M. Radman. Phenomenology of an inducible mutagenic dna repair pathway in escherichia coli: Sos repair hypothesis. In Molecular and Environmental Aspects of Mutagenesis. 1974.

J. M. Rendel. The canalization of the scute phenotype of Drosophila. Evolution, 13(4):425–439, June 1959.

J. M. Rendel. Bristle pattern in scute stocks of drosophila melanogaster. The American Naturalist, 99(904):25–32, 1965.

W. A. Rogers, S. Grover, S. J. Stringer, J. Parks, M. Rebeiz, and T. M. Williams. A survey of the trans-regulatory landscape for drosophila melanogaster abdominal pigmentation. Developmental biology, 385(2):417–432, 2014.

N. Rohner, D. F. Jarosz, J. E. Kowalko, M. Yoshizawa, W. R. Jeffery, R. L. Borowsky, S. Lindquist, and C. J. Tabin. Cryptic variation in morphological evolution: Hsp90 as a capacitor for loss of eyes in cavefish. Science, 342(6164):1372–1375, 2013.

S. L. Rutherford and S. Lindquist. Hsp90 as a capacitor for morphological evolution. Nature, 396: 336–342, 1998.

W. Scharloo. Canalization: genetic and developmental aspects. Annual Review of Ecology and Systematics, 22(1):65–93, 1991.

M. L. Siegal and A. Bergman. Waddington’s canalization revisited: Developmental stability and evolution. Proceedings of the National Academy of Sciences USA, 99:10528–10532, 2002.

George Gaylord Simpson. The baldwin effect. Evolution, 7(2):110–117, 1953.

Alexander J Stewart, Todd L Parsons, and Joshua B Plotkin. Environmental robustness and the adaptability of populations. Evolution, 66(5):1598–1612, 2012.

K. R. Thornton. A c++ template library for efficient forward-time population genetic simulation of large populations. Genetics, 198(1):157–166, 2014.

C. H. Waddington. Canalization of development and the inheritance of acquired characters. Nature, 150:563–565, 1942.

C. H. Waddington. Genetic assimilation of an acquired character. Evolution, 7(2):118–126, June 1953.

A. J. Wagers, J. L. Christensen, and I. L. Weissman. Cell fate determination from stem cells. Gene therapy, 9(10):606, 2002.

A. Wagner. Robustness and evolvability: a paradox resolved. Proceedings of the Royal Society of London B: Biological Sciences, 275(1630):91–100, 2008.

A. Wagner. The role of robustness in phenotypic adaptation and innovation. Proceedings of the Royal Society of London B: Biological Sciences, 279(1732):1249–1258, 2012.

A. Wagner. Does evolutionary plasticity evolve? Evolution, 50:1008–1023, 1996.

G.P. Wagner, G. Booth, and H. Bagheri-Chaichian. A population genetic theory of canalization. Evolution, 51:329–347, 1997.

Thomas Werner, Shigeyuki Koshikawa, Thomas M Williams, and Sean B Carroll. Generation of a novel wing colour pattern by the wingless morphogen. Nature, 464(7292):1143–1148, 2010.

M. R. Wheeler. Drosophilidae. In J. F. McAlpine, editor, Manual of Nearctic Diptera, volume 2, pages 1011–1018. Agriculture Canada, Research Branch Otawa, 1987.

